# A comparative and functional genomics analysis of the genus *Romboutsia* provides insight into adaptation to an intestinal lifestyle

**DOI:** 10.1101/845511

**Authors:** Jacoline Gerritsen, Bastian Hornung, Jarmo Ritari, Lars Paulin, Ger T. Rijkers, Peter J. Schaap, Willem M. de Vos, Hauke Smidt

## Abstract

Cultivation-independent surveys have shown that the recently described genus *Romboutsia* within the family *Peptostreptococcaceae* is more diverse than previously acknowledged. The majority of *Romboutsia*-associated 16S rRNA gene sequences have an intestinal origin, but the specific roles that *Romboutsia* species play in the digestive tract are largely unknown. The complete genomes of the human intestinal isolate *Romboutsia hominis* FRIFI^T^ (DSM 28814) and the soil isolate *Romboutsia lituseburensis* A25K^T^ (DSM 797) were sequenced. An evaluation of the common traits of this recently defined genus was done based on comparative genome analysis of the two strains together with the previously elucidated genome of the type species *Romboutsia ilealis* CRIB^T^. These analyses showed that the genus *Romboutsia* covers a broad range of metabolic capabilities with respect to carbohydrate utilization, fermentation of single amino acids, anaerobic respiration and metabolic end products. Main differences between strains were found in their abilities to utilize specific carbohydrates, to synthesize vitamins and other cofactors, and their nitrogen assimilation capabilities. In addition, differences were found with respect to bile metabolism and motility-related gene clusters.

## Introduction

The family *Peptostreptococcaceae* has recently been undergoing significant taxonomic changes. One of them involved the creation of the new genus *Romboutsia*, which currently contains the recognized species *Romboutsia ilealis, Romboutsia lituseburensis* (formerly known as *Clostridium lituseburense*) and *Romboutsia sedimentorum* (1, 2). Sequences related to the genus *Romboutsia* have been predominantly reported from samples of (mammalian) intestinal origin. Clone-library studies have shown the occurrence of *Romboutsia*-like 16S ribosomal RNA (rRNA) sequences in intestinal content samples (duodenum, jejunum, ileum and colon) from dogs (3) and cows (4), and in faecal samples from rats (5), polar bears (6) and porpoises (7). In addition, more recent sequencing-based studies have identified similar phylotypes in faecal samples from humans and other mammals (8), human ileal biopsies (9), mouse faecal samples (10), ileal biopsies from pigs (11), and ileal contents from rats (12) and deer (13). The isolation of strains from intestinal sources that possibly belong to other novel species within the genus *Romboutsia* has been reported as well. For example, strain TC1 was isolated from the hide of a cow, where it was found likely as the result of a faecal contamination of the hide (14). Furthermore, *Romboutsia timonensis* was isolated from human colon (15), and ‘*Clostridium dakarense’* was isolated from the stool of a 4-month-old Senegalese child (16). Recently, our search for a *Romboutsia* isolate of human small intestinal origin has resulted in the isolation of strain FRIFI from ileostoma effluent of a human adult and the subsequent proposal of *Romboutsia hominis* sp. nov. (17).

Although these findings suggest that members of the *Romboutsia* genus are mainly gut inhabitants, *Romboutsia* strains have been isolated from other environmental sources as well (2, 18, 19). The type strain of the second validly described species within the genus *Romboutsia, R. lituseburensis* A25K^T^, is not of intestinal origin, but was originally isolated from soil and humus from Côte d’Ivoire (20). Based on 16S rRNA gene identity, strains very similar to *R. lituseburensis* A25K^T^ have been isolated in recent years: strain H17 was isolated from the main anaerobic digester of a biogas plant (GI: EU887828.1), strain VKM B-2279 was isolated from a p-toluene sulfonate degrading community (21), strain 2ER371.1 was isolated from waste of biogas plants (22), and strain E2 was isolated from a cellulose degrading community enriched from mangrove soil (23). Furthermore, the type strain of the recently described novel species *R. sedimentorum* has been isolated from sediment samples taken from an alkaline-saline lake located in Daqing oilfield.

Altogether these studies suggest that the genus *Romboutsia* is probably more diverse than previously appreciated, and it is the question whether intestinal strains have adapted to a life outside a host or whether strains originating from other, non-host associated environments have adapted to a life in the intestinal tract. Because of the still limited availability of cultured representatives and their genomes, we know little about the specific roles that members of the genus *Romboutsia* play in the ecosystems in which they are found. To gain more insight into the metabolic and functional capabilities of the genus *Romboutsia*, we present here the genomes of the human intestinal isolate *R. hominis* FRIFI and the soil isolate *R. lituseburensis* A25K^T^, together with an evaluation of the common traits of this recently defined genus based on comparative genome analysis, including a comparison with the previously elucidated genome of the rat small intestinal isolate and type species of the genus, *R. ilealis* CRIB^T^.

## Materials and Methods

### Growth conditions and genomic DNA preparation

For genomic DNA extraction, *R. hominis* FRIFI^T^ and *R. lituseburensis* A25K^T^ (DSM 797) were grown overnight at 37°C in liquid CRIB-medium (pH 7.0) (1). DNA extraction was performed as described previously (24). DNA quality and concentrations were determined by spectrophotometric analysis using NanoDrop (Thermo Scientific) and by electrophoresis on a 1.0 % (w/v) agarose gel. DNA was stored at 4°C until subsequent sequencing.

### Genome sequencing and assembly

Genome sequencing of *R. hominis* FRIFI^T^ was carried out at the University of Helsinki (Finland) on a PacBio RS II, resulting in 134.366 PacBio reads and a total amount of 464.930.600 bases. Assembly was performed with PacBio SMRT analysis pipeline v2.2 and the HGAP protocol (25). Default settings were used except for: minimum sub-read length 500, minimum polymerase read length quality 0.80, minimum seed read length 7000, split target into chunks 1, alignment candidate per chunk 24, genome Size 3,000,000, target coverage 30, overlapper error rate 0.06, overlapper mini length 40, overlapper K-mer 14.

Genome sequencing of *R. lituseburensis* A25K^T^ was performed at GATC Biotech (Konstanz, Germany). One MiSeq library was generated on an Illumina MiSeq Personal Sequencer with 250 nt paired-end reads and an insert size of 500 nt, which resulted in 772.051 paired-end reads. Additionally one PacBio library was generated on a PacBio RS II, which resulted in 441.151 subreads and in total 998.181.178 bases. A hybrid assembly was carried out with MiSeq paired-end and PacBio CCS reads. For the MiSeq paired-end reads first all rRNA reads were removed with SortMeRNA v1.9 (26) using all included databases. Next, adapters were trimmed with Cutadapt v1.2.1 (27) using default settings except for an increased error value of 20 % for the adapters, and also using the reverse complement of the adapters. Quality trimming was performed with PRINSEQ Lite v0.20.0 (28) with a minimum sequence length of 40 and a minimum quality of 30 on both ends and as mean quality of the read. The assembly was done in parallel using two different assemblers. Ray v2.3 was used for the MiSeq paired-end dataset and the PacBio CCs dataset, using default settings except for a k-mer value of 75. The PacBio SMRT analysis pipeline v2.2 was run on the SMRT-cell subreads with the protocol RS_HGAP_Assembly_2, using default settings except for that the number of seed read chunks was set to 1, minimum seed read length was set to 7000, alignment candidate per chunk was set to 24 and the estimated genome size was reduced to 4 Mb. Both assemblies were merged, and duplications were identified based on BLASTN hits. Duplicate contigs were discarded if they had a hit with at least 99 % sequence identity within a bigger contig, which spanned at least 98 % of contig query length. Furthermore contigs with a length of less than 500 bp were discarded. The remaining contigs were merged with CAP3 v.12/21/07 (29), with an overlap length cut-off of 5000 bp and a minimum identity of 90 %. A circular element was detected within this assembly, based on BLASTP results of the predicted proteins (e-value 0.0001), and this was excluded from the further assembly process, but added to the final assembly result. Scaffolding of the contigs was done with SSPACE-LongRead (30) and the PacBio CCS reads using default settings. Further scaffolding was done with Contiguator v2.7.4 (31) using the genome of *R. hominis* FRIFI^T^ as reference genome and applying default settings. Copy numbers of the 16S rRNA gene from published genomes were derived from the rrnDB v4.0.0 (32). Average nucleotide identity (ANI) values were calculated with JSpecies v1.2.1 (33) by pairwise comparisons of available genomes within the family *Peptostreptococcaceae*.

### Genome annotation

Annotation was carried out with an in-house pipeline (adapted from (34)), with Prodigal v2.5 for prediction of protein coding DNA sequences (CDS) (35), InterProScan 5RC7 for protein annotation (36), tRNAscan-SE v1.3.1 for prediction of tRNAs (37) and RNAmmer v1.2 for the prediction of rRNAs (38). Additional protein function predictions were derived via BLAST searches against the UniRef50 (39) and Swissprot (40) databases (download August 2013). Subsequently the annotation was further enhanced by adding EC numbers via PRIAM version March 06, 2013 (41). Non-coding RNAs were identified using rfam_scan.pl v1.04, on release 11.0 of the RFAM database (42). COGs (43) were determined via best bidirectional blast (44), with an e-value of 0.0001. CRISPRs were annotated using CRISPR Recognition Tool v1.1 (45). A further step of automatic curation was performed, by weighting the annotation of the different associated domains, and penalizing uninformative functions (e.g. “Domain of unknown function”), and prioritizing functions of interest (e.g. domains containing “virus”, “phage”, “integrase” for phage related elements; similar procedure for different other functions).

Homology between the CDS of the *Romboutsia* strains was determined via best bidirectional BLAST hit (44) at the amino acid level with an e-value cut-off of 0.0001. To evaluate the core and pan metabolism of the *Romboutsia* strains, the three annotated genomes were supplied to Pathway tools v18 (46), and a limited amount of manual curation was performed to remove obvious false positives. Next the pathway databases were exported via the built-in lisp interface and the exported data was merged. A reaction was considered to be in the core metabolism if it was present in all three databases, else it was considered to be in the pan metabolism. Both parts were then reimported separately and combined into Pathway tools for further analyses.

Genes were matched to the list of essential and non-essential sporulation-related genes compiled by Galperin *et al.* (47) via different methods. Firstly, the protein-coding genes *of Bacillus subtilis* subsp. s*ubtilis* 168 were annotated via InterProScan and the respective *B. subtilis* sporulation-related proteins were matched to the proteins encoded by the three *Romboutsia* genomes, if they contained at least 50 % of the same domains. In case multiple matches were possible, the match with the highest domain similarity was picked. The matches were manually curated, and arbitrary proteins and/or false hits were excluded. For every protein, which did not have any match via the domains, the best bidirectional BLAST hit (e-value cut-off of 0.0001) was used instead. Secondly, the genome of *R. ilealis* CRIB^T^ was manually curated with respect to putative sporulation-related genes. In case the genomes of the other *Romboutsia* strains did not have any corresponding match for one of the proteins whereas a manually curated hit was present in *R. ilealis* CRIB^T^, the best bidirectional hit was assigned. Genomes were manually checked for further missing essential sporulation- and germination-related genes as defined by Galperin *et al.* (47). Function curation was performed with assistance of the *B. subtilis* wiki (http://subtiwiki.uni-goettingen.de/).

## Results and Discussion

To gain more insight in the metabolic and functional capabilities of members of the genus *Romboutsia* within an intestinal environment, we set out to elucidate the genome of a *Romboutsia* strain of human intestinal origin. To this end, the genome of *R. hominis* FRIFI^T^, isolated from ileostoma effluent of a human adult, was sequenced (17). For comparative analysis, we also aimed to determine the genome sequence of an isolate from another habitat, and thus the soil isolate *R. lituseburensis* A25K^T^ was obtained from the German Collection of Microorganisms and Cell Cultures (DSMZ, Braunschweig, Germany). Here we present the genome sequences of both organisms, together with an evaluation of the common traits of this recently defined genus based on comparative genome analysis, including the recently elucidated genome of the type species *R. ilealis* CRIB^T^ (1, 48). The genomes of *R. hominis FRIFI*^T^ and *R. lituseburensis* A25K^T^ (raw data and annotated assembly) have been deposited at the European Nucleotide Archive under project numbers PRJEB7106 and PRJEB7306, respectively.

Both *Romboutsia hominis* FRIFI^T^ (DSM 28814) and *R. lituseburensis* A25K^T^ (DSM 797) are anaerobic, Gram-positive, motile rods, belonging to the genus *Romboutsia. R. lituseburensis* A25K^T^ possesses the ability to sporulate. Pictures of both organisms can be seen in (Fig. 1).

**Fig. 1.**
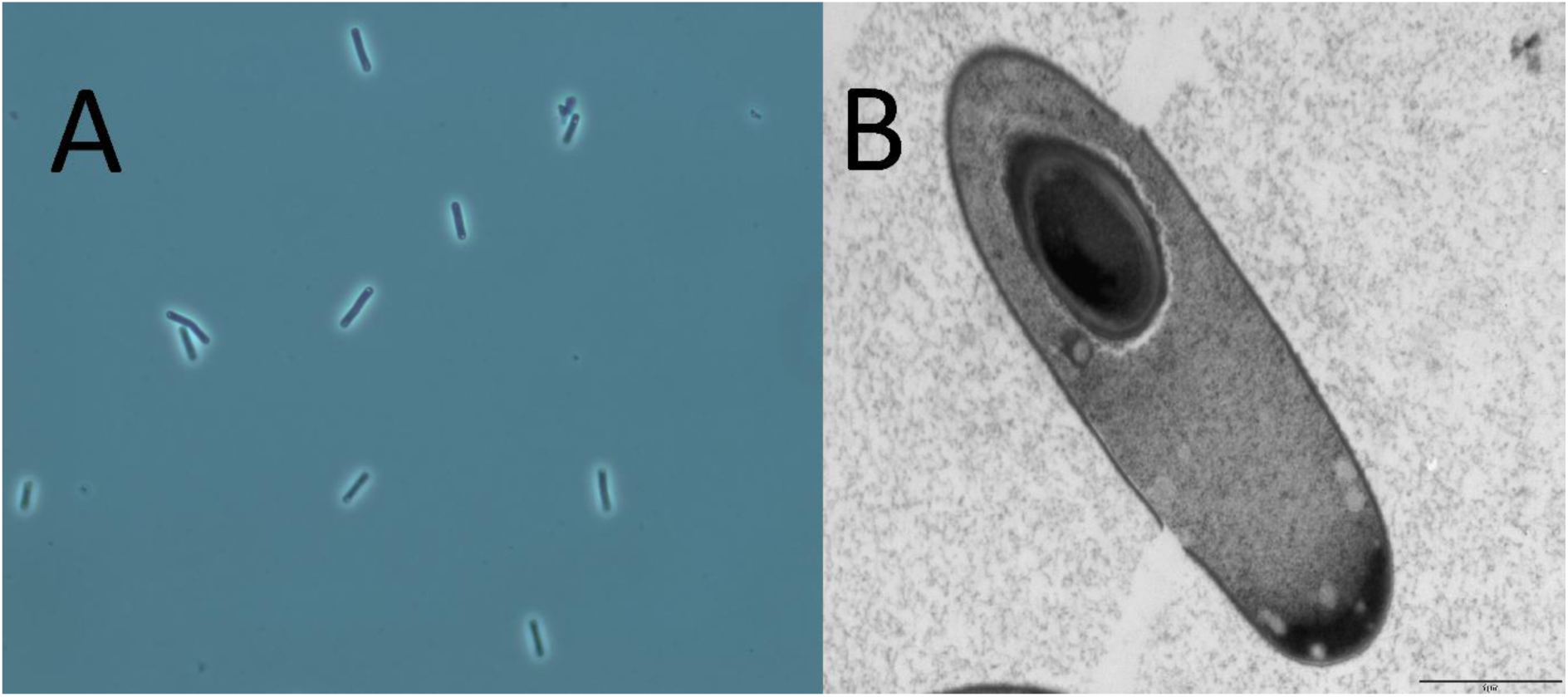
Electron micrographs of both Romboutsia species. A) Micrograph of *Romboutsia hominis* FRIFI^T^ B) Micrograph of *Romboutsia lituseburensis* A25K^T^.

To investigate the relationships between these isolates and their closest relatives, a 16S rRNA gene based neighbour joining-tree was constructed with a representative copy of the 16S rRNA gene of the type strains of the three species *R. ilealis, R. lituseburensis and R. hominis* (Fig. 2). Based on their 16S rRNA gene sequence these three species, together with the recently characterized species *Romboutsia sedimentorum*, form a monophyletic group.

**Fig. 2.**
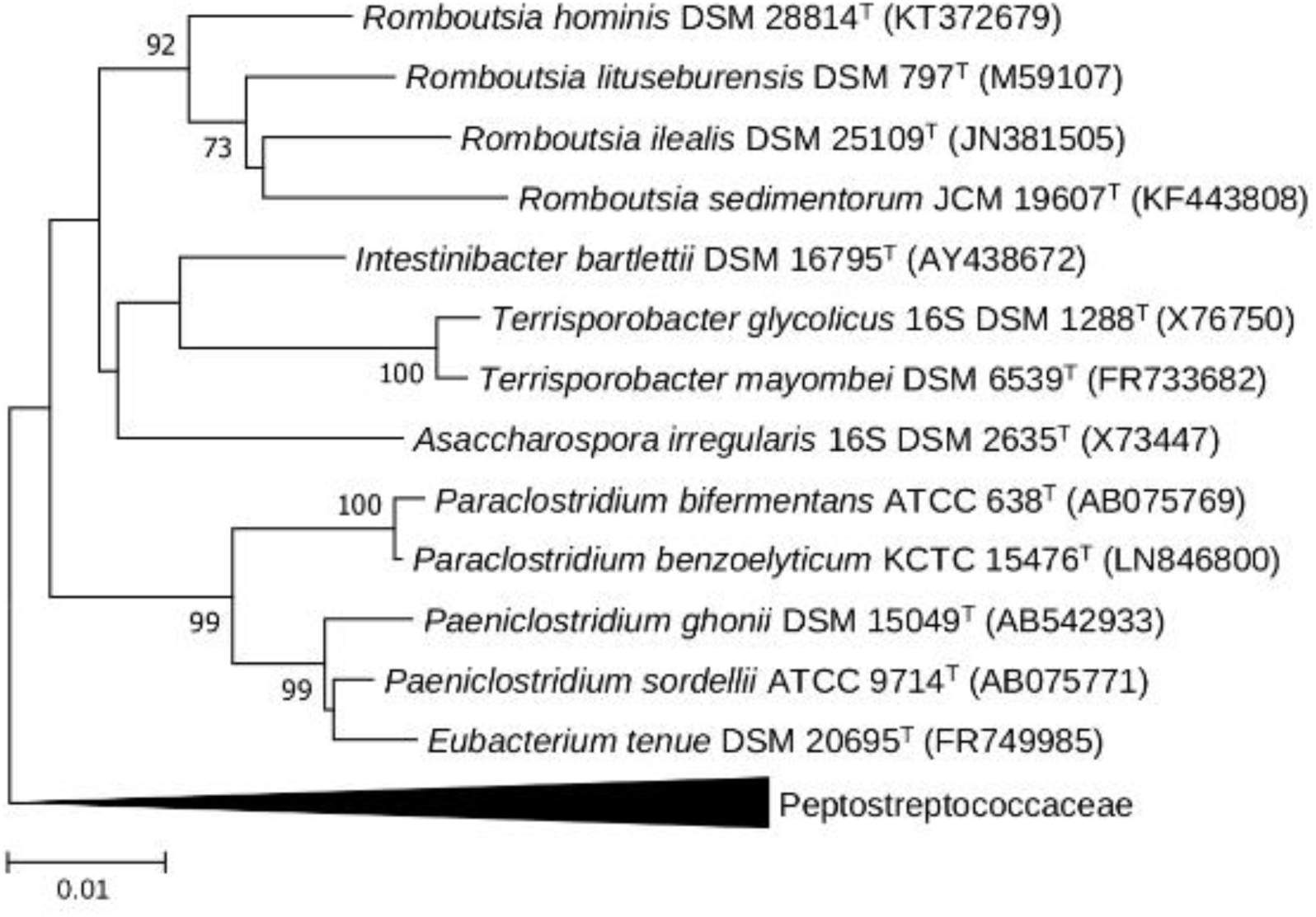
Neighbour-joining tree based on 16S rRNA gene sequences of *Romboutsia* species and closely-related species. The 16S rRNA gene sequences were aligned using the SINA aligner (49). The tree was constructed using MEGA6 software (50) with Kimura’s two-parameter model as substitution model. Only bootstrap values >70 % are shown at branch nodes. Bootstrap values were calculated based on 1000 replications. The reference bar indicates 1 % sequence divergence. GenBank accession numbers are given in parentheses. “The tree was rooted using 16S rRNA gene sequences of type strains of more distantly-related species within the family *Peptostreptococcaceae*.

*R. hominis* FRIFI^T^ contains a single, circular chromosome of 3.090.335 bp with an overall G+C content of 27.8 % (Table 1). The chromosome encodes 2.852 predicted coding sequences (CDS), of which 83 % have a function assigned. *R. lituseburensis* strain A25K^T^ contains one circular chromosome of 3.776.615 bp and one circular plasmid of 97.957 bp, with an overall G+C content of 28.2 % (Table 1). The chromosome encodes 3.535 CDS and the plasmid 123 CDS, of which 82 % have a function assigned. In addition, one segment of 4.101 bp, containing one 5S rRNA and four CDS (one two-component system and two subunits of an ABC transporter), could not be placed. The plasmid of *R. lituseburensis* A25K^T^ encodes two plasmid replication proteins, transporters, transcription factors, hydrolases and acyltransferases.

**Table 1.** General features of the *Romboutsia* genomes.

|  | *Romboutsia* sp.<br>strain FRIFI | *R. lituseburensis*<br>A25K<sup>T</sup> |
| --- | --- | --- |
| Status current assembly<br>of the chromosome | 1 scaffold,<br>1 gap | 2 scaffolds,<br>11 gaps |
| Genome size (Mb) | 3.09 | 3.88 |
| Chromosome size (Mb) | 3.09 | 3.78 |
| Plasmid size (Mb) | - | 0.98 |
| G+C content (%) | 27.8 | 28.2 |
| Total no. of CDS | 2852 | 3662 |
| No. of rRNA genes |  |  |
| 16S rRNA genes | 16 | --* |
| 23S rRNA genes | 17 | --* |
| 5S rRNA genes | 15 | --* |
| No. of tRNAs | 107 | 118 |
| No. of ncRNAs | 82 | 116 |
| CRISPR repeats | - | - |

\* Number of rRNA genes cannot accurately be estimated since some of the rRNA genes are situated next to an assembly gap of unknown size and might therefore be duplicates (*R. lituseburensis* A25K<sup>T</sup>)

The numbers of genes associated with general COG functional categories are shown in Table 2. The biggest differences between both genomes were found within the genes not assigned to any COG category. Within the COG categories, the most noticeable difference was observed within category J, with more than 1% difference. Most other categories are present in comparable abundances, despite the fact that both organisms were isolated from different habitats.

**Table 2.**
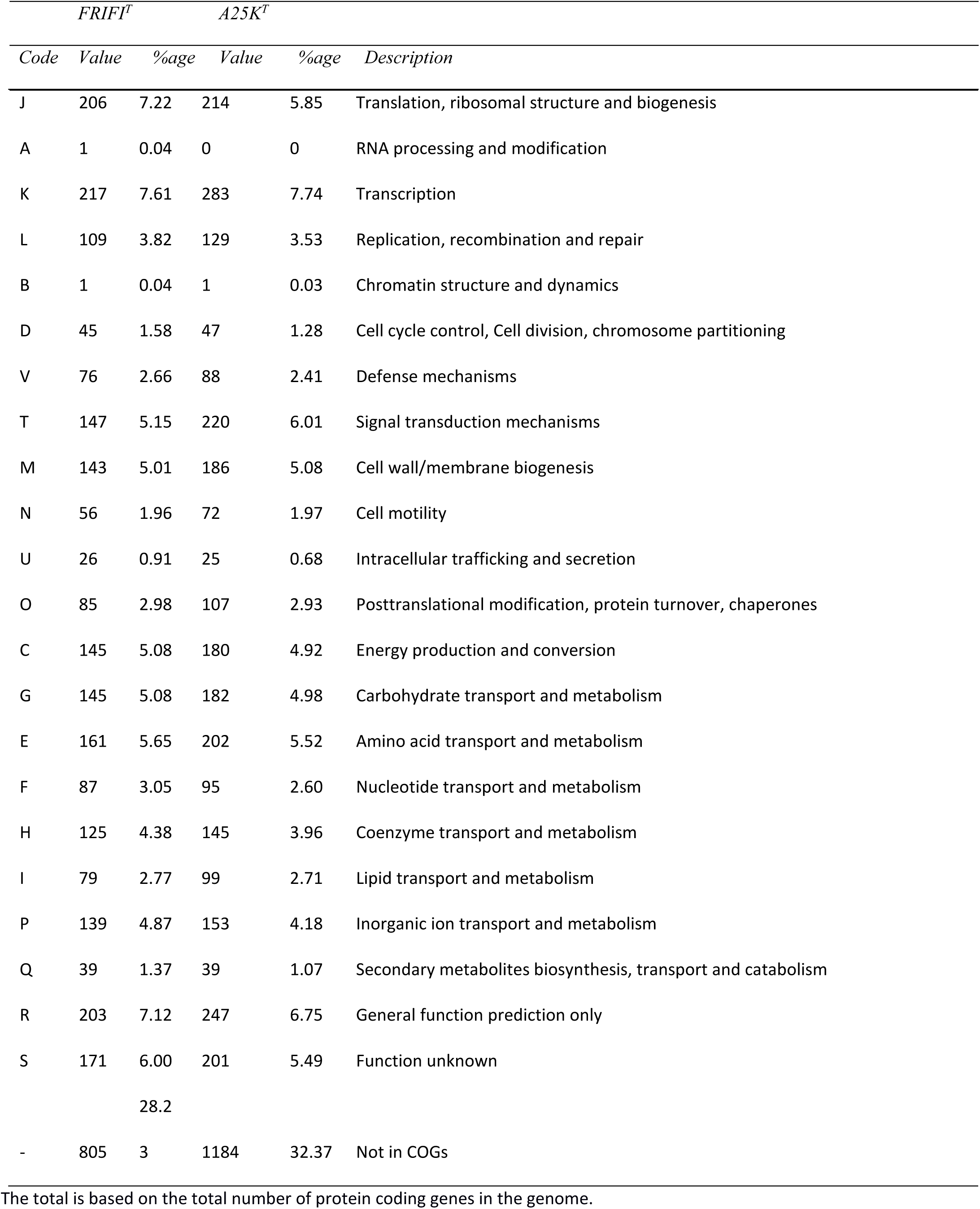
Number of genes associated with general COG functional categories.

### Impact of high number of ribosomal operons on sequencing efforts

Gaps in whole genome assemblies are usually located in repetitive regions that include ribosomal operons, which can appear multiple times in the genome. Also for the *Romboutsia* genomes, the presence of a high number of rRNA operons has been problematic for genome assembly. The assembly of *R. hominis* FRIFI^T^ contains one gap situated in a long stretch of ribosomal operons. The assembly of the chromosome of *R. lituseburensis* A25K^T^ contains eleven gaps, of which six are generated due to scaffolding with the use of a reference. Nine of the eleven gaps are situated within or neighbouring rRNA operons or tRNA clusters. A total of 16 copies of the 16S rRNA gene were identified in the genome of *R. hominis* FRIFI^T^. This is one of the highest copy numbers reported for the 16S rRNA gene in prokaryotic species up to this date. For *R. lituseburensis* A25K^T^ the total number of 16S rRNA genes could not be accurately estimated since some of the rRNA genes are situated next to assembly gaps, but at least 15 rRNA operons seem to be present. Pairwise sequence identity of the 16S rRNA sequences showed that within the genome of strain FRIFI^T^ there was an average sequence identity of 99.3 % and the lowest identity between individual copies was 98.4 %. Sequence divergence in the 16S rRNA gene is not uncommon within individual prokaryotic genomes (51, 52). However, for *R. hominis* FRIFI^T^ the divergence is located in the V1-V2 region of the 16S rRNA gene, one of the regions that is commonly used for sequence-based bacterial community analyses (53). In this region the average sequence identity was only 96.5 % and the lowest identity was only 92.3 %. Consequence of this divergence is that during identity clustering in operational taxonomic units (OTUs) the different copies of the 16S rRNA gene of *R. hominis* FRIFI^T^ end up in different OTUs, even at the level of 97 % identity, resulting in an overestimation of the diversity in *Romboutsia* phylotypes. In comparison, the type species of the genus, *R. ilealis* CRIB^T^, contains little variation in the 16S rRNA gene sequence (>99 % sequence identity), despite that also in this genome 14 copies of the 16S rRNA gene were identified.

### Comparison of the genome of *Romboutsia lituseburensis* to other genomes of this species

The genome sequence of *R. lituseburensis* A25K^T^ was compared to the genome sequence of the same strain (Bioproject PRJEB16174) that has been sequenced by the JGI and that has become publicly available during the course of this project (54). This comparison showed only minor sequence differences. Both genome sequences, including the plasmid sequences, are nearly identical (99.9%), with most differences arising from the gaps within our assembly or from contig ends (∼500bp of each contig) within the JGI assembly. One difference was observed within the repetitive gene RLITU_1618, which was shorter assembled in the JGI genome. The surface antigen encoding gene RLITU_0237 was also assembled shorter within the JGI assembly. Duplicated Lysine, Serine and Arginine tRNA genes (location 1433719 - 1434325) were omitted in the JGI assembly, probably due to misassembly in this complicated region, which is not resolvable with Illumina short reads. Furthermore the JGI contig GI: 1086420641 seems to be assembled differently, since the rRNA region present in this contig was not connected to the protein coding sequences in our assembly, but both were located at different places within the genome. We were unable to locate the first 8kb of JGI contig GI: 1086420759 within our assembly. The remaining parts of this contig match to an area following an assembly gap, and it therefore cannot be excluded that it was missed in our assembly. The only unplaced contig within our assembly was also nearly completely contained in a single contig within the JGI assembly, and therefore did not help to resolve this situation. Overall, it seems that most of the observed differences were due to technical reasons, and not due to underlying differences in the genomes of both strains.

### Comparative genomic analysis of the new *Romboutsia* genomes to the type species of the genus

The genome sequences of the two newly sequenced strains were compared to the type strain of the type species of the genus, *Romboutsia ilealis* CRIB^T^ (48). The number of protein coding genes per genome within the various strains was quite variable, ranging from 2359 to 3658 (Table 1). The number of putative homologous genes among the three *Romboutsia* genomes was determined via amino acid level best bidirectional hits (Fig. 3). In total 1522 genes were shared between all three strains, the core genome, accounting for 42 % to 64 % of the total gene count in the individual genomes, providing a first insight in the genomic heterogeneity within the genus. The bigger the genome, the more unique genes were present, ranging from 19 % to 34 % of the total gene count.

**Fig. 3.**
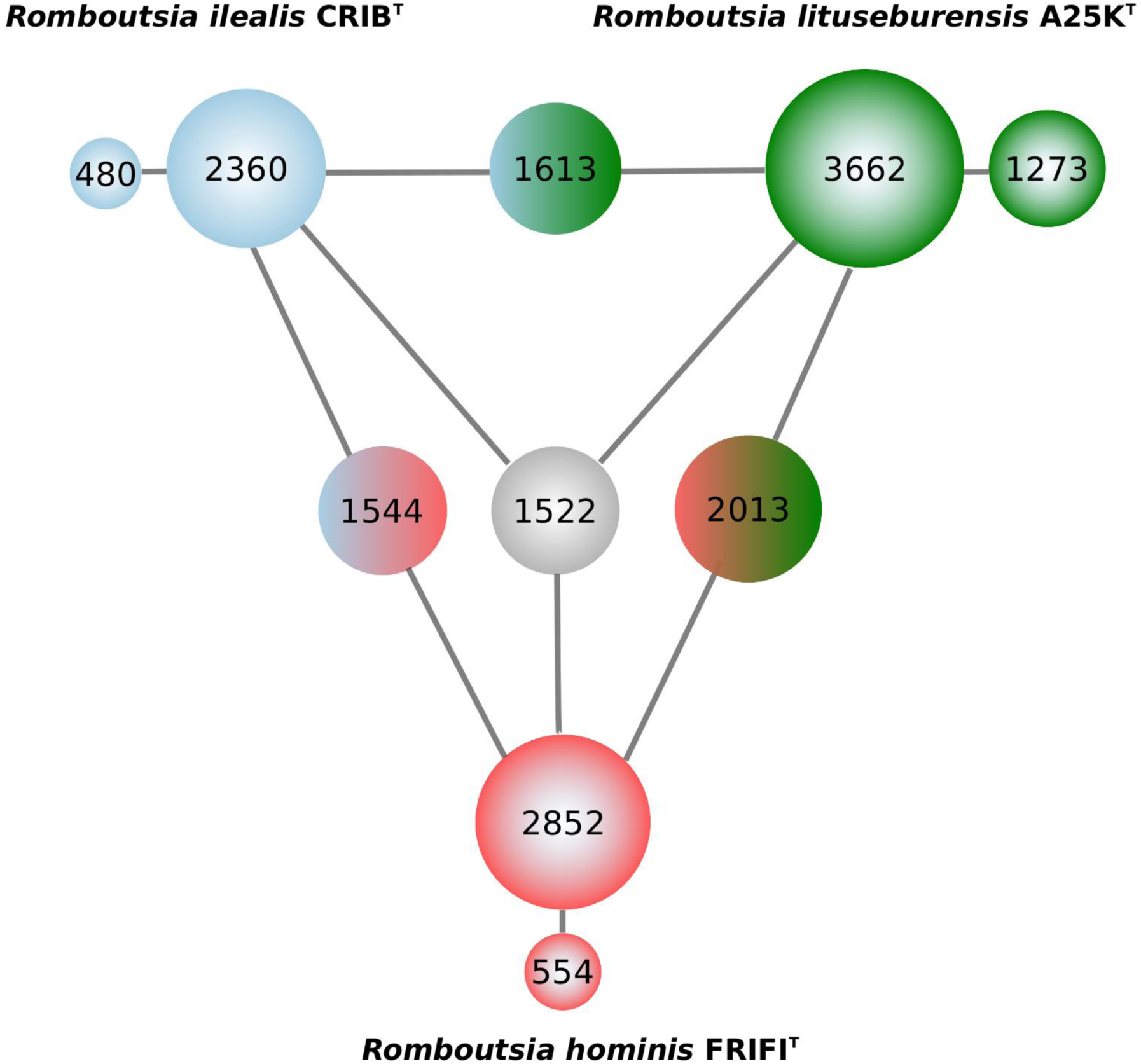
Overview of the number of homologous genes shared between the three *Romboutsia* genomes. The circles are colour-coded by the *Romboutsia* strains they represent: blue, *R. ilealis* CRIB^T^ green; *R. lituseburensis* A25K^T^; red *R. hominis* FRIFI^T^. Also the total number of genes and the number of unique genes are indicated for each genome. The area of the circle is representative for the number of genes.

The comparative genome analysis showed a general conservation of the genomic structure of the genus *Romboutsia* around the replication start site, while synteny appears to be lost towards the replication end point. For most pairwise comparisons, synteny was lost at a quarter of the genome in both up- and downstream directions, making roughly half of the genomes syntenic. Breaks of synteny appear to be related to specific recombination events. For example, compared to the other genomes synteny is absent in *R. ilealis* CRIB^T^ due to the insertion of a prophage, whereas the regions both up- and downstream are syntenic. At another spot in the genome of *R. ilealis* CRIB^T^ synteny is lost due to phage-related genes found around the tmRNA gene, which has been reported to be a common insertion site for phages (55). The position of the tmRNA itself is roughly equal in all three genomes, but no synteny could be observed in its vicinity. Strain/species-specific gene clusters, like the CRISPR-Cas system or the fucose degradation pathway present in *R. ilealis* CRIB^T^, appear to be situated more towards the less conserved replication end point. One point of conservation in the less conserved area is an inversion of one part of the butyrate fermentation pathway, which is absent in *R. ilealis* CRIB^T^, but inverted between *R. hominis* FRIFI^T^ and *R. lituseburensis* A25K^T^. Some significant deletion events appear to have occurred, since they can be observed in the conserved areas of the genomes. For example, the pili encoding gene cluster, which is found in the genome of *R. lituseburensis* A25K^T^ close to the replication start site, is absent in the genomes of *R. ilealis* CRIB^T^ and *R. hominis* FRIFI^T^ except for a twitching motility protein encoding gene. Another example is the biosynthesis cluster for vitamin B12, which is also located in all strains close to the replication start site. While this cluster is complete in *R. lituseburensis* A25K^T^ and *R. hominis* FRIFI^T^, only remnants of the cluster are visible in *R. ilealis* CRIB^T^ as there is a deletion of nine genes, which prevents the biosynthesis of cob(II)yrinate a,c-diamide. This cluster is also situated next to an rRNA operon, of which only the one in *R. ilealis* CRIB^T^ has an integrase inserted.

### Core and pan metabolism of the genus *Romboutsia*

An overview of the core metabolism of the *Romboutsia* strains and strain-specific metabolic features is provided in Fig. 4. All three *Romboutsia* strains can ferment carbohydrates via the glycolysis, and possess the non-oxidative pentose phosphate pathway. Moreover, from the genomes it was predicted that all strains have the capability to synthesize (and degrade) all nucleotides, cell wall components, fatty acids and siroheme. In addition, it was predicted that all three *Romboutsia* spp. strains can only produce a limited non-identical set of amino acids. In turn they are, however, also able to ferment numerous amino acids. Furthermore various pathways for assimilation of nitrogen were predicted, as well as a pathway for production of the quorum sensing compound autoinducer AI-2. Some of the metabolic highlights will be discussed in the following paragraphs.

**Fig. 4.**
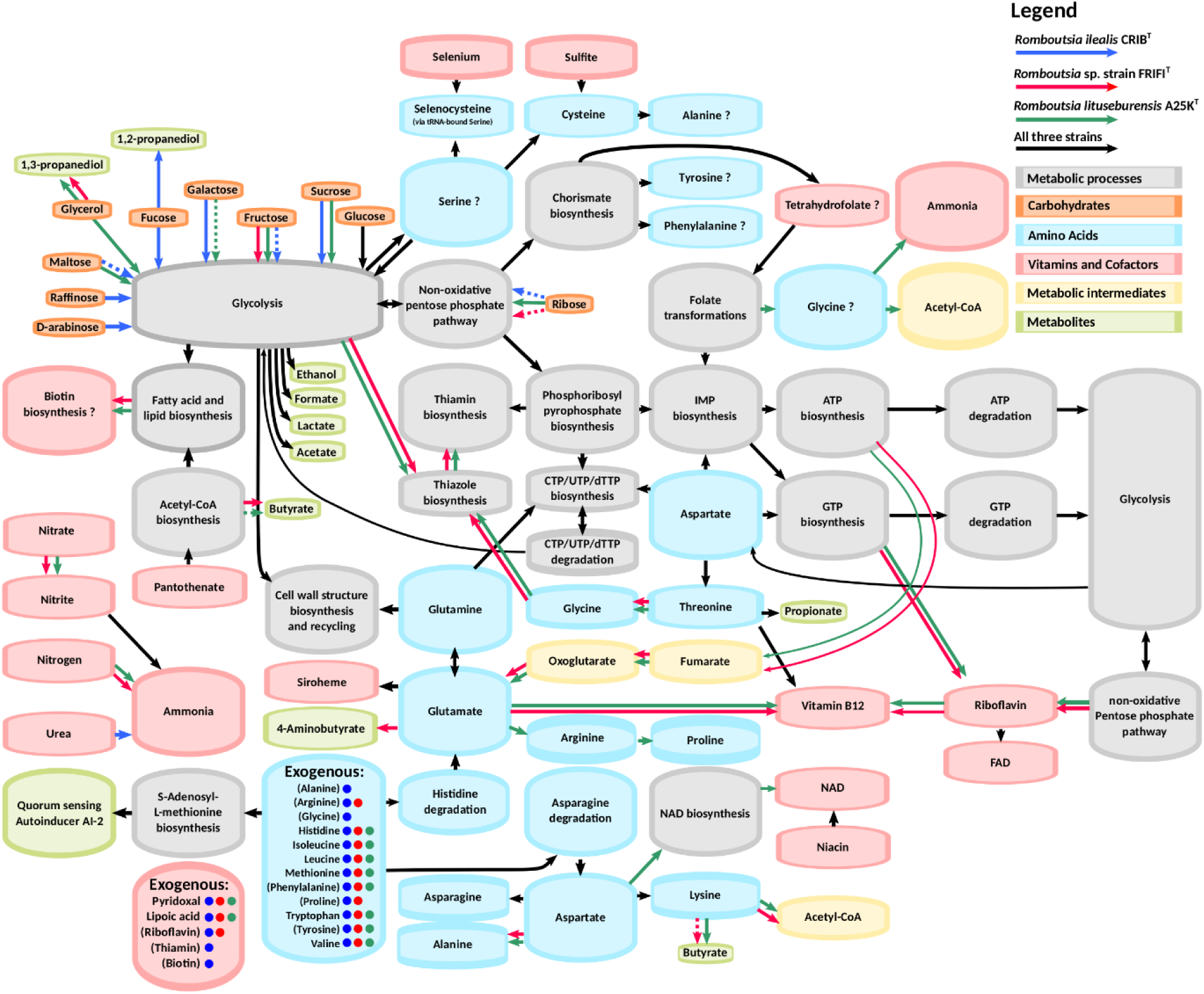
Overview of the predicted core metabolism and strain-specific metabolic features of three *Romboutsia* spp. strains. Dotted lines indicated instances where reported experimental observations do not match genome-based predictions. Question marks indicate processes for which not all enzymes could be identified in the genome. Brackets around compounds indicate that the enzymes necessary for de novo production of the compounds might be present in the genome.

### Fermentation and anaerobic respiration

Similar fermentation end-products have been observed for *R. ilealis* CRIB^T^, *R. hominis* FRIFI^T^ and *R. lituseburensis* A25K^T^ during growth on glucose, including formate, acetate and a small amount of lactate (17). The pathways leading to formate, acetate and lactate production, which have previously been described for *R. ilealis* CRIB^T^ (48), were also found in the two other *Romboutsia* strains suggesting that all strains are indeed able to produce formate, acetate and lactate. Butyrate (and iso-valerate) production has been observed for *R. lituseburensis* A25K^T^ during *in vitro* growth on undefined medium components such as beef extract, peptone and casitone (but not on yeast extract). The addition of a carbohydrate (e.g. glucose) resulted in a redirection of the fermentation pathways towards other end products such as formate (data not shown). Two pathways leading to butyrate synthesis, the acetyl-CoA and the lysine pathways, could be predicted from the genome of *R. lituseburensis* A25K^T^. The pathways are co-located in the genome, suggesting that the acetoacetyl-CoA formed during lysine fermentation can be directly used as substrate in the acetyl-CoA pathway for additional energy conservation (56, 57). A lysine-specific permease was predicted in the genome as well, suggesting that exogenous lysine can serve as energy source for this strain. Since an acetyl-CoA acetyltransferase was found in the gene cluster as well, a fully functioning carbohydrate-driven acetyl-CoA pathway is expected. For the final step in butyrate production, a phosphate butyryl transferase/butyrate kinase (*buk*) gene cluster was identified in the genome. Similar gene clusters (although with some gene inversions) were found in the genome of *R. hominis* FRIFI^T^ as well, but butyrate production has not (yet) been observed (17).

In the genomes of both *R. hominis* FRIFI^T^ and *R. lituseburensis* A25K^T^ a reductive pathway for the metabolism of glycerol was predicted, comprising a glycerol dehydratase and 1,3-propanediol dehydrogenase (58). This suggests that these strains are able to ferment glycerol and produce 1,3-propanediol as one of the fermentation end-products. Production of 1,3-propanediol has indeed been reported for *R. lituseburensis* (22). Furthermore, in the genome of *R. lituseburensis* A25K^T^ the oxidative pathway for glycerol metabolism, including glycerol dehydrogenase and dihydroxyacetone kinase, could be identified as well, suggesting that this strain should be able to use glycerol as sole carbon and energy source. For both *R. hominis* FRIFI^T^ and *R. lituseburensis* A25K^T^ growth on glycerol has indeed been observed (17), although the responsible genes could not be predicted in *R. hominis* FRIFI^T^.

The genomes of all three *Romboutsia* strains studied here contain genes encoding for enzymes of the Wood-Ljungdahl pathway. A formate dehydrogenase was predicted for all strains except *R. ilealis* CRIB^T^. The presence of formate dehydrogenase together with a complete Wood-Ljungdahl pathway categorizes them as potential acetogens, microbes that can grow autotrophically using CO_2_ and H_2_ as carbon and energy source. This provides them with metabolic flexibility in addition to heterotrophic growth on organic compounds. The role of acetogens in the intestinal tract is not well studied. They have been proposed to play an important role in hydrogen disposal, in addition to methanogens and sulfate reducers (59, 60).

Genomes of all three *Romboutsia* strains contain genes predicted to encode a sulfite reductase of the AsrABC type. Inducible sulfite reductases are directly linked to the regeneration of NAD^+^, which plays a role in energy conservation and growth, as well as to detoxification of sulfite (61). *R. hominis* FRIFI^T^, however, appears to lack the formate/nitrite transporter family protein that was found in the vicinity of the predicted sulfite reductase in the other strains similarly to *Clostridioides difficile* (previously known as *Clostridium difficile*) where it was characterized as a hydrosulfide ion channel which exports the toxic metabolites from the cell (62). The genes coding for a complete membrane-bound electron transport system were identified in both genomes of *R. hominis* FRIFI^T^ and *R. lituseburensis* A25K^T^, similar to the Rnf system identified in microbes such as *Clostridium tetani, Clostridium ljungdahlii* and *C. difficile.* In these species the system is suggested to be used to generate a proton gradient for energy conservation in microbes without cytochromes. In *C. tetani*, the system is proposed to play a role in the electron flow from reduced ferredoxin, via NADH to the NADH-consuming dehydrogenase of the butyrate synthesis pathway (63). In addition, the Rnf system is proposed to be used by *C. ljungdahlii* during autotrophic growth (64). In the genome of *R. ilealis* CRIB^T^only remnants of an Rnf electron transport system could be found, which might be a result of genome reduction since also no complete butyrate synthesis pathway or acetogenic pathways were found.

### Fermentation of individual amino acids

Species belonging to the class *Clostridia* are known for their capabilities to ferment amino acids. Of the three *Romboutsia* strains, *R. lituseburensis* A25K^T^ appears to be the most resourceful. All three *Romboutsia* strains are predicted to be able to ferment L**-**histidine via glutamate using a histidine ammonia lyase. In addition, fermentation of L-threonine was predicted using a L**-**threonine dehydratase resulting in propionate production, which has been described for *R. lituseburensis* (20). Fermentation of L-serine into pyruvate using an L-serine dehydratase was predicted for all three strains as well. As aforementioned, *R. hominis* FRIFI^T^ and *R. lituseburensis* A25K^T^ are predicted to be able to ferment L**-**lysine. In addition, *R. lituseburensis* A25K^T^ is predicted to ferment glycine using the glycine reductase pathway found in other related species including *C. difficile* (65, 66). A corresponding complex has also been identified in *R. hominis* FRIFI^T^, but it is likely to be non-functional, due to a loss of two of the three subunits. Furthermore, the ability to ferment L**-**arginine (using an arginine deiminase) and L**-**glutamate (using a Na^+^-dependent glutaconyl-CoA decarboxylase) was predicted for *C. lituseburensis* A25K^T^ as well. A glutamate decarboxylase was predicted for *R. hominis* FRIFI^T^, suggesting the ability to decarboxylate glutamate to 4-aminobutyrate (GABA) for this strain only.

### Amino acid and vitamin requirements

Pathways for (*de novo*) synthesis of amino acids were identified in the three *Romboutsia* strains (Table 3). All three strains show similar dependencies on exogenous amino acid sources. Based on genome predictions, *R. lituseburensis* A25K^T^ is able to synthesize lysine from aspartate, whereas the last enzyme in this pathway is missing in the genomes of *R. hominis* FRIFI^T^ and *R. ilealis* CRIB^T^. In addition, *R. hominis* FRIFI^T^ and *R. lituseburensis* A25K^T^are predicted to synthesise alanine from aspartate and glycine from threonine. Common to all organisms is that the prephenate dehydratase for the biosynthesis of phenylalanine and tyrosine is missing, although all other enzymes for the biosynthesis of chorismate and for the further conversion to both amino acids are present.

**Table 3.** Overview of genome-based predictions for amino acid requirements of the three *Romboutsia* strains. In case only one or two enzymes are missing in either salvage or *de novo* pathway leading to the production of an amino acid, this is indicated in parentheses.

|  | *Romboutsia*<br>*hominis* FRIFl<sup>T</sup> | *R. lituseburensis*<br>*A25K*<sup>T</sup> | *R. ilealis*<br>*CRIB*<sup>T</sup> |
| --- | --- | --- | --- |
| **Alanine** | + | + | - |
| **Arginine** | - | + | - |
| **Asparagine** | + | + | + |
| **Aspartate** | + | + | + |
| **Cysteine** | + | + | + |
| **Glutamate** | + | + | + |
| **Glutamine** | + | + | + |
| **Glycine** | + | + | - |
| **Histidine** | - | - | - |
| **Isoleucine** | - | - | - |
| **Leucine** | - | - | - |
| **Lysine** | - (-1) | + | - (-1) |
| **Methionine** | - | - | - |
| **Phenylalanine** | - (-1) | - (-1) | - (-1) |
| **Proline** | - | - (-1) | - |
| **Serine** | - (-1) | - (-1) | - (-1) |
| **Threonine** | - (-1) | - (-1) | - (-1) |
| **Tryptophan** | - | - | - |
| **Tyrosine** | - (-1) | - (-1) | - (-1) |
| **Valine** | - | - | - |

The urease gene cluster, previously identified in *R. ilealis* CRIB^T^ (48), could not be identified in the two other *Romboutsia* strains. However, a nitrogenase encoding gene cluster was identified in the genomes of *R. hominis* FRIFI^T^ and *R. lituseburensis* A25K^T^, suggesting that these two strains are able to fix N_2_.

All three strains encode one or several oligopeptide transporters of the OPT family (67). In addition, two oligopeptide transport systems (*Opp* and *App*) (68, 69) were predicted in *R. hominis* FRIFI^T^ and *R. lituseburensis* A25K^T^ (strain FRIFI^T^ misses the OppA), whereas they were absent in *R. ilealis* CRIB^T^. Based on the predicted amino acid dependencies, it can be concluded that these *Romboutsia* strains are adapted to an environment rich in amino acids and peptides.

The metabolic capabilities of the three *Romboutsia* species are comparable regarding the ability to produce certain vitamins and other cofactors (Fig. 4). None of them is predicted to be able to synthesize vitamin B6, lipoic acid or pantothenate, but it is likely that they are all able to produce siroheme from glutamate and CoA from pantothenate. As previously described for *R. ilealis* CRIB^T^ (48), the pathway for *de novo* folate biosynthesis via the pABA branch is present, however, a gene encoding dihydrofolate reductase could not be identified in any of the three *Romboutsia* strains. However, since this enzyme is essential in both *de novo* and salvage pathways of tetrahydrofolate, it is highly likely it is present in the genomes. The biosynthetic capabilities of *R. lituseburensis* A25K^T^, and *R. hominis* FRIFI^T^ are larger than that of *R. ilealis* CRIB^T^, as they are both predicted to produce biotin, thiamin and vitamin B12. The gene clusters for biotin and thiamin biosynthesis are located in the more variable regions of the genomes as discussed above, and the vitamin B12 biosynthesis pathway is incomplete in *R. ilealis* CRIB^T^ due to a deletion, as mentioned earlier. Only *R. lituseburensis* A25K^T^ is predicted to have the capacity to produce riboflavin *de novo.* Furthermore, *R. lituseburensis* A25K^T^ is, as the only non-host derived organism in this comparison, the only strain that can synthesize NAD *de novo*.

### Bile resistance

One of the challenges for microbes living in the intestinal tract is that they have to deal with the host-secreted bile acids. The bile acid pool size and composition modulates the size and composition of the intestinal microbiota and vice versa (70, 71). Bile acids can undergo a variety of bacterial transformations including deconjugation, dehydroxylation and epimerization. In both intestinal isolates, *R. ilealis* CRIB^T^ and *R. hominis* FRIFI^T^, a choloylglycine hydrolase encoding gene was identified. Bile salt hydrolases (BSHs), also known as conjugated bile acid hydrolases, from the choloylglycine hydrolase family are widespread among Gram-positive and Gram-negative intestinal microbes (72). They are involved in the hydrolysis of the amide linkage in conjugated bile acids, releasing primary bile acids. There is a large heterogeneity among BSHs, including with respect to their substrate specificity such as specificity towards either taurine or glycine conjugated bile salts (70). The choloylglycine hydrolases of *R. ilealis* CRIB^T^ and *R. hominis* FRIFI^T^ differ significantly from each other (32 % identity at amino acid level), suggesting a different origin. The BSH of *R. ilealis* CRIB^T^ and *R. hominis* FRIFI^T^ show at the amino acid level 52 % and 33 % identity, respectively, with the choloylglycine hydrolase CBAH-1 from *Clostridium perfringens* (73).

In addition to the possible BSH, a bile acid 7α-dehydratase encoding gene could be identified in *R. hominis* FRIFI^T^. This enzyme is part of the multi-step 7α/ß-dehydroxylation pathway that is involved in the transformation of primary bile acids into secondary bile acids. So far, this pathway has been found exclusively in a small number of anaerobic intestinal bacteria all belonging to the *Firmicutes* (72, 74). The presence of this pathway enables microbes to use primary bile acids as an electron acceptor, allowing for increased ATP formation and growth. High levels of secondary bile acids are associated with diseases of the host such as cholesterol gallstone disease and cancers of the GI tract (75, 76). However, the evidence that bacteria capable of 7α-dehydroxylation are directly involved in the pathogenesis of these diseases is still limited. The pathway has been extensively studied in the human isolate *Clostridium scindens* VPI 12708 (formerly known as *Eubacterium* sp. strain VPI 12708 (77)). In addition, 7α-dehydroxylation activity was also reported for *Clostridium hiranonis* (78) and *Paeniclostridium sordellii* (previously known as *Clostridium sordellii*) (79), and other close relatives of the genus *Romboutsia* (74). Extensive characterization of the 7α/ß-dehydroxylation pathway in *C. scindens* VPI 12708 has demonstrated that the genes involved are encoded by a large bile acid inducible (*bai*) operon (72). For *R. hominis* FRIFI^T^ several other genes were identified in the vicinity of the bile acid 7α-dehydratase gene that showed homology to the genes in the *bai* operon, but some other (essential) genes seem to be missing. From gene presence/absence it was therefore not possible to predict whether *R. hominis* FRIFI^T^ has 7α/ß-dehydroxylation activity and that has to be confirmed experimentally.

### Toxins and virulence-related genes

The class *Clostridia* contains some well-known pathogens, including *C. difficile, C. botulinum* and *C. perfringens*, for which several toxins have been characterized in depth (80). No homologues of the genes coding for the toxins of *C. difficile* (toxin A, toxin B, binary toxin) or *C. botulinum* could be found in the genomes of the three *Romboutsia* strains. The genome of *R. ilealis* CRIB^T^ encodes a predicted protein that was annotated as a putative septicolysin (CRIB_2392) since it shares 56 %identity to a protein that has been characterized as an oxygen-labile hemolysin in *Clostridium septicum* (81). However, the exact role of septicolysin in potential pathogenesis is not known. Homologues are not found in other related species. A homologue for the alpha toxin of *Clostridium perfringens* (80, 82), a phospholipase C protein involved in the aetiology of gas gangrene caused by *C. perfringens* (83), was found by BLAST search in the genomes of *R. hominis* FRIFI^T^ and *R. lituseburensis* A25K^T^ (49.4 - 54.3 % identity at the amino acid level). In addition, *R. lituseburensis* A25K^T^ is predicted to contain a protein homologous to the perfringolysin O (theta toxin) of *C. perfringens*, which is a thiol-activated cytolysin that forms large homo-oligomeric pore complexes in cholesterol-containing membranes, which is also involved in gas gangrene aetiology. By BLAST search similar proteins could also be found in the genomes of *P. sordellii* and *Paraclostridium bifermentans* (previously known as *Clostridium bifermentans*), which are close relatives of the genus *Romboutsia*. There are many homologous enzymes produced by other bacteria that do not have similar toxigenic properties as the *C. perfringens* proteins (83). For example, the phospholipase C proteins produced by *P. bifermentans* and *P. sordellii* were found to have significantly less haemolytic activity than the homologuous protein of *C. perfringens* (51 and 53.4 % similarity on amino acid level, respectively) (84, 85). The predictions for the presence of potential toxin-encoding genes in the *Romboutsia* strains was done based on homology; the enzymatic activity of the gene products will have to be determined in the future to see whether some of the *Romboutsia* strains have toxigenic properties.

### Motility

Motility was observed for *R. hominis* FRIFI^T^ and *R. lituseburensis* A25K^T^ but not for *R. ilealis* CRIB^T^, as previously reported (17). In general, different appendages can be found on bacterial surfaces that provide bacteria with the ability to swim in liquids or move on surfaces via gliding or twitching motility (86, 87). In the genomes of *R. hominis* FRIFI^T^ and *R. lituseburensis* A25K^T^ a large gene cluster for the synthesis of flagella could be identified. The organization of the flagella gene cluster is very similar to that in the genome of *C. difficile*. The formation of flagella involves a whole array of different components, including the core protein flagellin. Post-translational modification of flagellin by glycosylation is an important process both for the flagellar assembly and biological function, and genes involved in these modifications are often found in the vicinity of the structural flagellin genes (88). This is also the case for *R. hominis* FRIFI^T^ and *R. lituseburensis* A25K^T^, and these genes are found in an intra-flagellar synthesis locus similar to the situation in *C. difficile* 630 (89). In *R. ilealis* CRIB^T^ no genes encoding flagellar proteins or genes involved in chemotaxis could be identified in line with the lack of motility (90). Flagella are dominant innate immune activators in the intestinal tract as flagellin molecules can be recognized by host cell-surface and cytoplasmatic pattern recognition receptors (91-93). The role for flagella in virulence and pathogenicity of *C. difficile* is a topic of interest, however, their exact contribution is still unknown (94). The flagellin proteins of some of the most abundant motile commensal microbes that are found in the human intestinal tract (*Eubacterium* and *Roseburia* species) have recently also been shown to possess pro-inflammatory properties (95).

One type of gliding motility involves the extension, attachment and retraction of type IV pili (TFP), which pull the bacterium towards the site of attachment (96). In the genome of *R. lituseburensis* A25K^T^ a complete set of genes for the assembly of Type IV pili could be identified. In contrast, in the genomes of *R. ilealis* CRIB^T^ and *R. hominis* FRIFI^T^ only remnants could be identified.

### Sporulation

Sporulation is a trait found only in certain low G+C Gram-positive bacteria belonging to the *Firmicutes* (97). The formation of metabolically dormant endospores is an important strategy used by bacteria to survive environmental challenges such as nutrient limitation. These endospores are resistant to extreme exposures (e.g. high temperatures, freezing, radiation and agents such as antibiotics and most detergents) that would kill vegetative cells. The ability to form endospores was also studied for the three *Romboutsia* strains. *R. lituseburensis* A25K^T^ readily forms mature spores, especially during growth in Duncan-Strong medium and Cooked meat medium, that both contain large quantities of proteose peptone, and spore formation was observed in almost every cell (data not shown). Previously, the endospore forming capabilities of *R. ilealis* CRIB^T^ and *R. hominis* FRIFI^T^ have been studied (90). Using different media and incubation conditions it was observed that the process of sporulation appears to be initiated, however, no free mature spores could be observed.

The whole process of sporulation and subsequent spore germination involves the expression of hundreds of genes in a highly regulated manner. At a molecular level the process is best understood in the model organism *Bacillus subtilis* (98). For species belonging to the class *Clostridia*, the process of sporulation is mainly studied in microbes in which sporulation has been shown to play a big role in other processes such as virulence (*C. perfringens, C. difficile, C. botulinum* and *C. tetani*) or solvent production (*Clostridium acetobutylicum*). Studying these microbes has made it clear that there are significant differences in the sporulation and germination process in species belonging to the class *Clostridia* compared to members of the *Bacilli* (47, 99, 100). The *B. subtilis* proteins involved in the early stages of sporulation, i.e. onset (stage I), commitment and asymmetric cell division (stage II), and engulfment (stage III), are largely conserved in *Clostridia* species. However, many of the proteins that play a role in later stages, i.e. cortex formation (stage IV), spore coat maturation (stage V), mother cell lysis and spore release (stage VII), appear to be less conserved. For example, limited spore outer layer conservation was observed in *C. difficile* compared to *B. subtilis* (101). Comparative genomic based-studies have tried to define the minimal set of genes essential for sporulation in clostridial species, but this has appeared to be challenging (47, 100, 102). In all spore-formers, initiation of sporulation is controlled by the transcription factor Spo0A, a highly conserved master regulator of sporulation. Phosphorylation of Spo0A leads to the activation of a tightly regulated cascade involving several sigma factors that regulate the further expression of a multitude of genes involved in sporulation. There are, however, significant differences in the regulation of the sporulation pathway between different clostridial species (103) of which we do not completely understand the impact on sporulation itself, highlighting that there is still a big gap in our knowledge on the complex process of sporulation.

The genomes of the three *Romboutsia* genomes were mined for homologues of sporulation specific genes according to the publication of Galperin *et al*. (47). All three *Romboutsia* strains have similar sets of sporulation-related genes, with *R. ilealis* CRIB^T^ having the least number of genes (147 genes) and *R. lituseburensis* A25K^T^ having the most (183 genes) (**Table S1**). The only protein that is deemed essential for sporulation, but which was only found in the genome of *R. lituseburensis* A25K^T^, is the Stage V sporulation protein S which has been implicated to increase sporulation (104). For *R. lituseburensis* A25K^T^, the sporulation regulator Spo0E was predicted to be absent, due to a point mutation in the start codon of the corresponding gene, changing it to an alternative start codon. This regulator is suggested to be involved in the prevention of sporulation under certain circumstances (105); impact of the point mutation on the presence/absence of the protein and on regulation of sporulation in *R. lituseburensis* A25K^T^ will have to be determined. Interestingly, the stage V sporulation proteins AA and AB, encoded by *spoVAA* and s*poVAB*, that are essential for sporulation in *Bacilli* since mutants lead to the production of immature spores (106), are absent in sporulating *Clostridium* species, but are present in all three *Romboutsia* strains. Furthermore, *R. lituseburensis* A25K^T^ is the only strain that contains the *sps* operon that has been shown to be involved in spore surface adhesion (107). Absence of this operon in *B. subtilis* resulted in defective germination, and more hydrophobic and adhesive spores, however, given that these proteins are also absent in nearly all clostridial species, their role in sporulation and germination in the *Romboutsia* strains still has to be determined. As also noted by Galperin *et al. (47)*, there are other species that have been demonstrated to be spore-forming but which also lack some of the genes that are deemed to be essential, e.g. *spoIIB, spoIIM*, and other proteins from the second sporulation stage in *Lysinibacillus sphaericus* C3-41v (47). In comparison, it is interesting to note that the genome of *C. hiranonis*, a close relative of the genus Romboutsia (and *C. difficile)*, appears to contain only 21 of the essential sporulation genes, missing for example most of the proteins related to the second and third stage of sporulation, while *C. hiranonis* is known to form spores ((108), and own observations). Altogether, based on gene presence/absence it is not possible to predict whether these *Romboutsia* strains are indeed able to successfully complete the process of sporulation and release endospores. An asporogenous phenotype could be credited to the absence or mutation of a single gene.

Initiation of sporulation is still a topic of interest. Accessory gene regulatory (*agr)*-dependent quorum sensing, and thus most likely cell density, has been proposed to play an important role in efficient sporulation (109). For *C. difficile*, however, quorum-sensing has been shown not to play a role in initiation of sporulation, and recently a more direct link between nutrient availability and sporulation was suggested (110). The uptake of peptides by the Opp and App oligopeptide transport systems appears to prevent initiation of sporulation in nutrient rich environments (69). Both transport systems are absent in *R. ilealis* CRIB^T^, but are present in the two other *Romboutsia* strains.

During sporulation, a number of species produce inclusion bodies and granules that are visible by phase contrast and electron microscopy. This is also true for *R. lituseburensis* A25K^T^ in which electron translucent bodies are visible in TEM pictures (Fig. 1), similar to the carbohydrate or polyhydroxybutyrate inclusions observed in for example *Clostridium pasteurianum* (111), *C. acetobutylicum* (112) and *C. botulinum* (113). The development of these inclusion bodies appears to coincide with the initiation of sporulation. Based on this observation, it can be speculated that by intracellular accumulation of a carbon and energy source these microbes ensure they can complete the sporulation process with only limited dependence on external carbon and energy sources.

## Conclusions

Based on the comparative genome analysis presented here we can conclude that the investigated genomes of the genus *Romboutsia* encode a versatile array of metabolic capabilities with respect to carbohydrate utilization, fermentation of single amino acids, anaerobic respiration and metabolic end products. A relative genome reduction is observed in the isolates from intestinal origin. In addition, the presence of bile converting enzymes and pathways related to host-derived carbohydrates, point towards adaption to a life in the (small) intestine of mammalian hosts. For each *Romboutsia* strain unique properties were found. However, since currently only one genome was available for each species, it is impossible to unequivocally predict which properties might apply to each species and which properties are strain-specific. Isolation and genome sequencing of additional strains from diverse environments is needed to provide a more in-depth view of the metabolic capabilities at the species- as well as the genus level and to reveal specific properties that relate to adaptation to an intestinal lifestyle.

## Availability of data and material

All data has been uploaded to the European Nucleotide Archive under project numbers PRJEB7106 and PRJEB7306

## Authors’ contributions

Conceptualization: Hauke Smidt, Willem M. de Vos

Investigation: Jacoline Gerritsen, Bastian Hornung, Jarmo Ritari, Lars Paulin

Supervision: Hauke Smidt, Ger T. Rijkers, Peter J. Schaap

Writing – original draft preparation: Jacoline Gerritsen, Bastian Hornung, Hauke Smidt

Writing – Review & Editing: Jacoline Gerritsen, Bastian Hornung, Ger T. Rijkers, Hauke Smidt

## Acknowledgements

The authors would like to thank Wilma Akkermans-van Vliet for help with DNA extractions, Aleksander Umanetc for providing the microscopic picture of *R. hominis* FRIFI^T^, and Laura van Niftrik for providing the TEM picture of *R. lituseburensis* A25K^T^. We would also like to thank Jasper Koehorst for help with the annotation, Bart Nijsse for assisting with the different software packages, and Jesse van Dam for help with the Pathway Tools lisp interface. In addition, we would like to thank William Trimble from the Argonne National Laboratory for helpful discussions concerning genome assembly with MiSeq data sets.

## Supplementary information

### File #1

- File name: supplementary_table_1.xls
- File format including the correct file extension: Excel document, .xls
- Title of data: Supplementary Table 1
- Description of data: Overview of sporulation-related genes in the three *Romboutsia* genomes. Genes from *Bacillus subtilis* subsp. *subtilis* 168, to which no homologues could be identified, are omitted. In case multiple candidate loci were detected, all are mentioned. Loci which are assigned to more than one gene from *B. subtilis* subsp. *subtilis* 168 are marked with an asterisk

